# Dynamic functional connectivity changes in dementia with Lewy bodies and Alzheimer’s disease

**DOI:** 10.1101/374538

**Authors:** Julia Schumacher, Luis R. Peraza, Michael Firbank, Alan J. Thomas, Marcus Kaiser, Peter Gallagher, John T. O’Brien, Andrew M. Blamire, John-Paul Taylor

## Abstract

We studied the dynamic functional connectivity profile of dementia with Lewy bodies (DLB) and Alzheimer’s disease (AD) and the relationship between dynamic connectivity and the temporally transient symptoms of cognitive fluctuations and visual hallucinations in DLB.

Resting state fMRI data from 31 DLB, 29 AD, and 31 healthy control participants were analysed using dual regression to determine between-network functional connectivity. We used a sliding window approach followed by k-means clustering and dynamic network analyses to study dynamic functional connectivity changes associated with AD and DLB. Network measures that showed significant group differences were tested for correlations with clinical symptom severity.

AD and DLB patients spent more time than controls in sparse connectivity configurations with absence of strong positive and negative connections and a relative isolation of motor networks from other networks. Additionally, DLB patients spent less time in a more strongly connected state and the variability of global brain network efficiency was reduced in DLB compared to controls. However, there were no significant correlations between dynamic connectivity measures and clinical scores.

The loss of global efficiency variability in DLB might indicate the presence of an abnormally rigid brain network and the lack of economical dynamics, factors which could contribute to an inability to respond appropriately to situational demands. However, the absence of significant clinical correlations indicates that the severity of transient cognitive symptoms such as cognitive fluctuations and visual hallucinations might not be directly related to these dynamic connectivity changes observed during a short resting state scan.

## 1. Introduction

Resting state functional MRI has been used to study changes in functional connectivity associated with different forms of dementia such as dementia with Lewy bodies (DLB) and Alzheimer’s disease (AD) (Kenny et al., 2012; Lowther et al., 2014; Peraza et al., 2014; Schumacher et al., 2018). To date, most functional connectivity studies have focused on mean connectivity over the duration of a scan of several minutes, thereby implicitly assuming that functional connectivity remains stationary during that time. However, it has recently been shown that functional connectivity can vary substantially in both strength and directionality on a timescale of seconds to minutes (Chang and Glover, 2010; Hutchison et al., 2013b) and that studying these dynamics can provide important complementary information to the traditional analysis of stationary functional connectivity (Calhoun et al., 2014; Hutchison et al., 2013a). A large number of DLB patients experience fluctuations in cognition and attention/arousal, mostly spontaneously occurring without any situational explanation, which results in pronounced variation in cognitive ability over time (Bradshaw et al., 2004; McKeith et al., 2005).

In addition, the majority of DLB patients present with visual hallucinations that recur over time (Aarsland et al., 2001). The transient nature of these DLB symptoms suggests that changes in functional connectivity dynamics might play a role in their aetiology. We therefore studied dynamic functional connectivity in DLB compared to healthy controls and patients with AD to (1) identify the differential dynamic connectivity profile of the two dementia subtypes and (2) investigate how abnormal dynamics might relate to the clinical DLB symptoms, especially with respect to cognitive fluctuations and visual hallucinations.

## 2. Materials and methods

### 2.1 Participants

The study involved 102 participants over 60 years of age: 33 were diagnosed with probable DLB, 36 with probable AD, and 33 were age-matched healthy controls (HCs) with no history of psychiatric or neurological illness. Participants from two contemporary and independent dementia studies conducted at one research center (Newcastle) were combined for this analysis. Both studies recruited patients from the local community-dwelling population who had been referred to old age psychiatry and neurology services. DLB and AD diagnoses were performed independently by two experienced old age psychiatrists using consensus criteria for probable DLB (McKeith et al., 2005) and probable AD (McKhann et al., 1984, 2011). Written informed consent was obtained from all participants and both studies were approved by the local ethics committee.

### 2.2 Data acquisition

Imaging for both studies was performed on the same 3T Philips Intera Achieva scanner. The imaging protocol was the same in both studies except for a different resolution of the structural scans. Structural images were acquired with a magnetization prepared rapid gradient echo (MPRAGE) sequence, sagittal acquisition, echo time 4.6ms, repetition time 8.3ms, inversion time 1250ms, flip angle = 8°, SENSE factor = 2, and in-plane field of view 256 x 256 mm^2^ with slice thickness 1.2 mm, yielding a voxel size of 0.93 x 0.93 x 1.2 mm^3^ (study 1) and in-plane field of view 240 x 240 mm^2^ with slice thickness 1.0 mm, yielding a voxel size of 1.0 x 1.0 x 1.0 mm^3^ (study 2). Resting state scans were obtained with a gradient echo echo-planar imaging sequence with 25 contiguous axial slices, 128 volumes, anterior-posterior acquisition, in plane resolution = 2.0 x 2.0 mm, slice thickness = 6 mm, repetition time (TR) = 3000ms, echo time = 40ms, and field of view = 260 x 260 mm^2^. DLB patients who were taking dopaminergic medication were scanned in the ON state.

### 2.3 Preprocessing

FEAT (FMRI Expert Analysis Tool, version 6.0) which is part of the FMRIB’s software library (FSL, www.fmrib.ox.ac.uk/fsl) was used to perform motion correction with MCFLIRT, slice-timing correction, and spatial smoothing with a 6.0mm full width at half maximum Gaussian kernel. Participants were excluded if the MCFLIRT motion parameters exceeded 2 mm translation and/or 2° rotation. To ensure that there were no group differences in motion due to the presence of Parkinsonian symptoms in the DLB patients, motion was compared between the groups using the formula introduced in (Liao et al., 2010).

A denoising procedure was performed with ICA-AROMA in FSL which performs single-subject independent component analysis (ICA) to remove motion components from each participant’s functional data (Pruim et al., 2015). Subsequently, eroded CSF and white matter masks were estimated using FAST in FSL and the mean signal inside the mask was regressed out of each participant’s cleaned functional data. Functional and structural images were co-registered using boundary based registration in FSL, and normalized to standard MNI space using Advanced Normalization Tools (Avants et al., 2011). As a final step, functional data were temporally high-pass filtered with a cut-off of 150 s and resampled to a resolution of 4 x 4 x 4 mm^3^.

### 2.4 Analysis of resting state data

Resting state networks (RSNs) were estimated with an independent set of 42 healthy control participants from two previous studies that were conducted on the same MR scanner using a similar imaging protocol (see Supplementary Table S1). Data from all 42 HCs were temporally concatenated and subjected to a group-ICA using FSL’s MELODIC. A meta ICA approach was adopted to obtain more reliable components (Biswal et al., 2010) using a model order of 70 independent components which has been shown to be optimal for assessing disease-related group differences (Abou Elseoud et al., 2011) (see Schumacher *et al*. (2018) for a more detailed description). Meta ICA components were visually inspected with respect to their spatial maps (Kelly et al., 2010) and 27 were identified as being of biological interest according to previous literature (Agosta et al., 2012; Beckmann et al., 2005) (Figure 1 and Supplementary Table S2).

**Figure 1:**
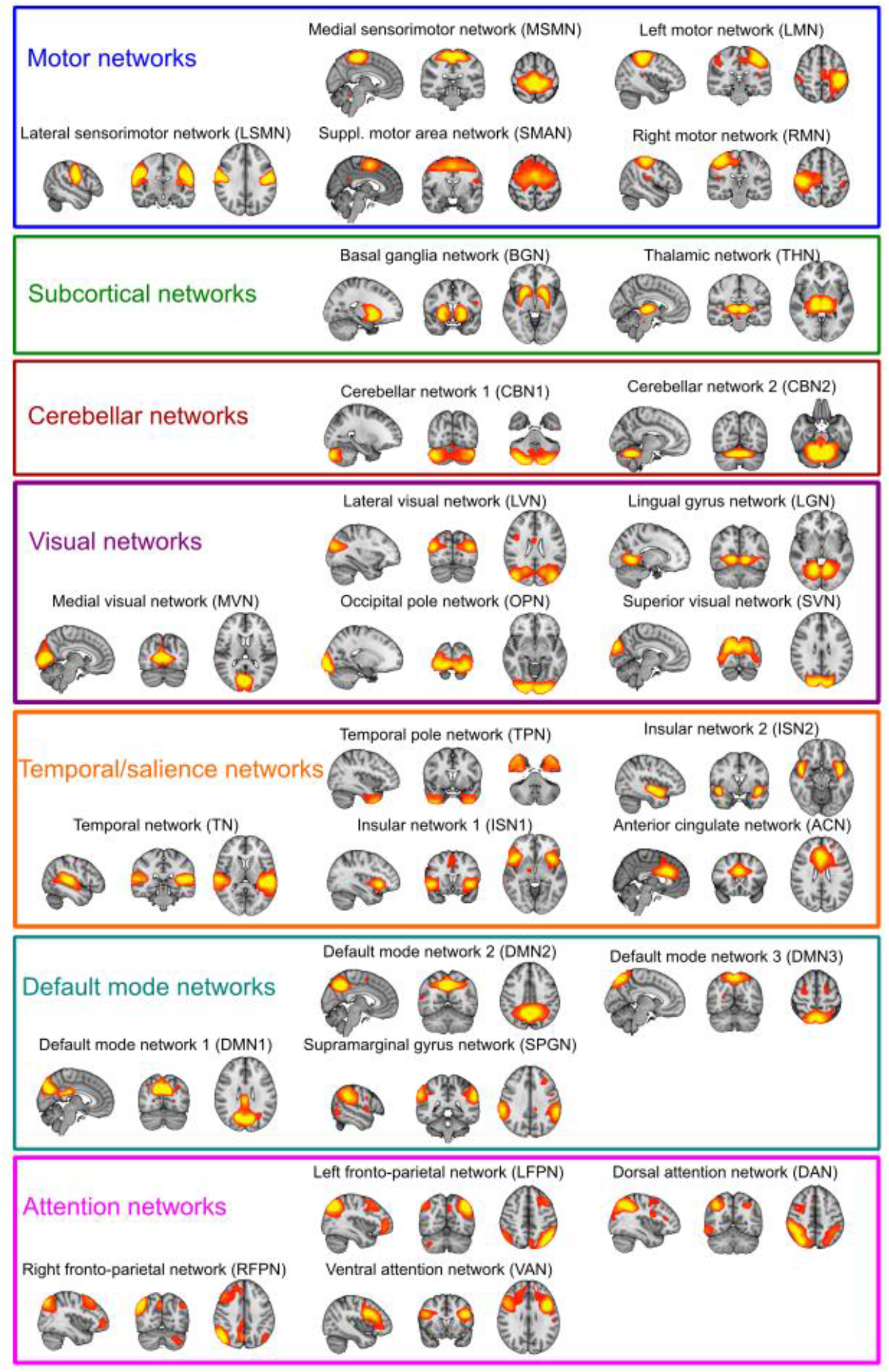
Resting state networks. Spatial maps of the 27 (RSNs) obtained from the independent healthy control group. RSN maps are thresholded at 3<z<12. Images are shown in radiological convention, i.e. the left side of the image corresponds to the right hemisphere.

Subsequently, FSL-dual regression was run with all 27 identified RSNs concatenated in a single 4D image, to obtain subject-specific representations of the RSN spatial maps and associated subject-specific time courses (Figure 2A). Results from a static connectivity analysis have been published previously using the same data (Schumacher et al., 2018).

**Figure 2:**
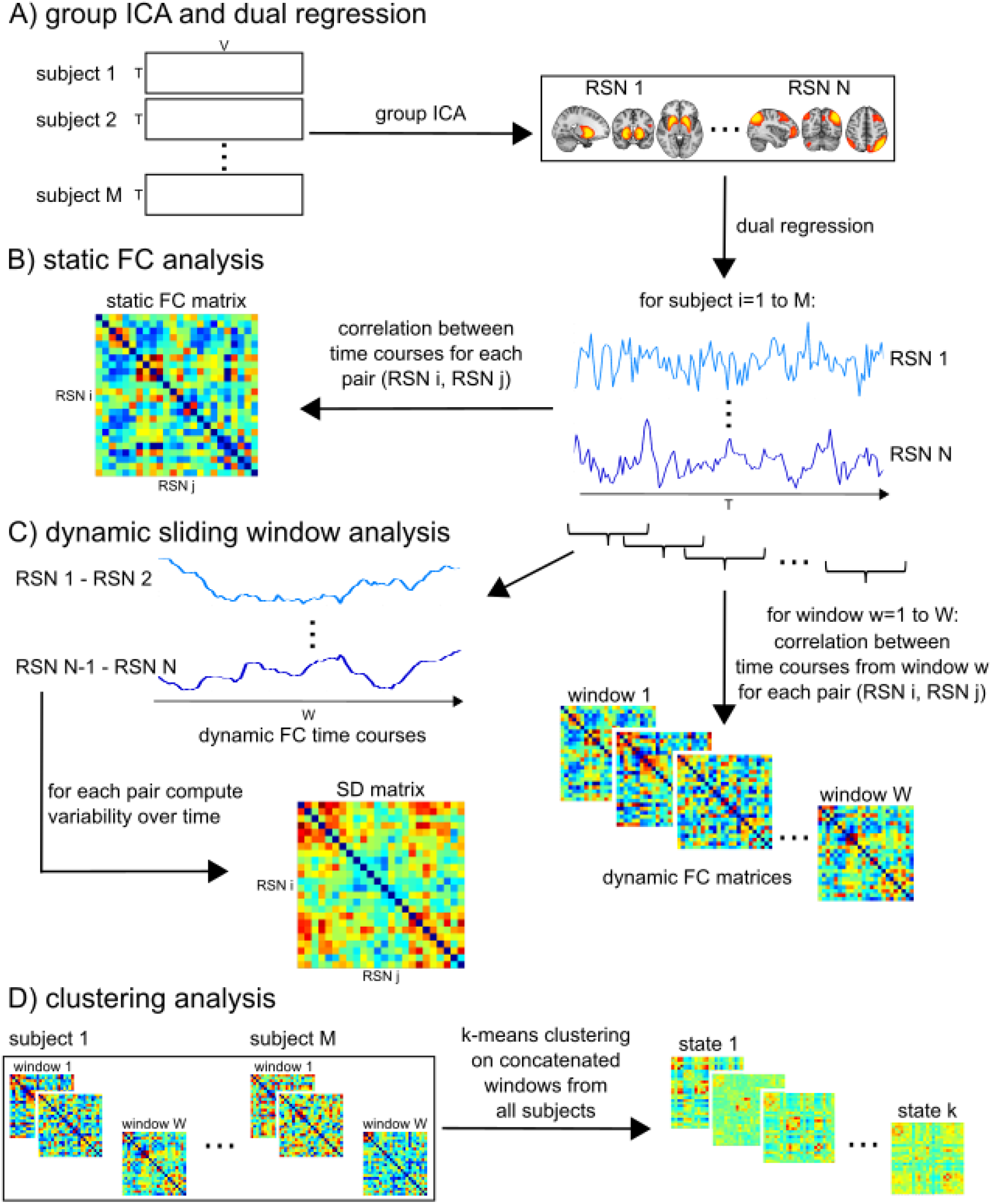
Sliding window approach and k-means analysis. A) Data from all healthy control subjects from the independent cohort is concatenated in time and subjected to group ICA to identify RSN spatial maps. Subject-specific time courses of each RSN are estimated using dual regression. B) static functional connectivity (FC) analysis by calculating correlation between each pair of RSNs using the whole time course (see (Schumacher et al., 2018)). C) sliding window approach and estimation of SD of connectivity over time. D) k-means clustering.

The subject-specific time courses resulting from dual regression were further processed in Matlab (R2016b) using functions from the Group ICA of fMRI toolbox (GIFT, http://mialab.mrn.org/software/gift/index.html) to remove remaining noise sources. Postprocessing included (1) detrending to remove linear, quadratic, and cubic trends, (2) outlier detection based on AFNI’s 3dDespike function (http://afni.nimh.nih.gov/afni) and interpolation of outliers using a third-order spline fit to the clean parts of the time courses, and (3) low-pass filtering using a fifth-order Butterworth filter with a cutoff frequency of 0.15 Hz.

### 2.5 Sliding window analysis

The postprocessed dual regression time series were analysed with a sliding window method to assess changes in between-network connectivity over time (Figure 2C). This analysis was performed in Matlab (R2016b) based on functions from GIFT (Allen et al., 2014). A tapered window was created by convolving a rectangle of 22TR (66s) with a Gaussian with sigma of 3TR and moved in steps of 1TR resulting in a total of 107 overlapping time windows. To assess the robustness of the results with respect to different window sizes, all analyses were repeated for window sizes ranging from 18 to 28 TR. A covariance matrix between all RSN-to-RSN pairs was estimated for each window separately. Since estimation of covariance based on short time series can be noisy, we estimated the regularised inverse covariance matrix using the graphical LASSO approach. An L1-norm constraint was imposed on the inverse covariance matrix to promote sparsity. The regularization parameter λ was optimized for each participant individually by evaluating the log-likelihood of unseen time windows from the same participant using 20-fold cross-validation. All covariances were subsequently converted to correlation values and transformed into z-scores using Fisher r-to-z transformation. To control for the effect of possible covariates the z-scores were then residualized with respect to age, gender, and study membership using multiple linear regression (Damaraju et al., 2014).

The variability of the connection strengths between RSNs (dynamic functional connectivity) was assessed by calculating the standard deviation (SD) of the RSN-to-RSN correlations across time windows. To assess whole-brain dynamics the mean SD across all connections between RSN pairs was computed. Additionally, the mean SD for each network across all other networks was considered and each RSN-to-RSN connection was also tested separately.

### 2.6. K-means clustering

To assess patterns of functional connectivity that reoccur over time across different participants, k-means clustering was applied to the windowed covariance matrices from all windows and all participants using the Manhattan (L1) distance function (Figure 2D). The optimal number of clusters k was chosen based on the elbow criterion of the cluster validity index, computed as the ratio of within-cluster to between-cluster distance (Allen et al., 2014). The clustering algorithm was repeated 500 times in Matlab with random initializations of cluster centroid positions to get a stable solution. In addition to using the optimal value for k, the analyses were repeated for k ranging from 2 to 8 to assess the robustness of the results regarding different values of k.

Group differences were assessed with respect to (1) frequency: proportion of windows assigned to a state, (2) mean dwell time: average number of consecutive windows assigned to a state, (3) intertransition interval: average number of consecutive windows before a state transition occurs, (4) number of transitions: overall number of transitions between different states (Hutchison and Morton, 2015; Marusak et al., 2016).

### 2.7. Dynamic network analysis

We also considered a graph-theoretic approach to study the dynamics of global and local efficiency using the Brain Connectivity Toolbox (Rubinov and Sporns, 2010). For each time window a graph was constructed using the 27 RSNs as nodes and the correlation between the RSNs within the respective time window as edge strength. We created binarized, unweighted, and undirected graphs by thresholding the absolute value of the individual time window correlation matrices to achieve different edge densities. The edge density of a graph is defined as the number of existing edges divided by the maximum number of possible edges (351 in our case). We used edge density thresholds ranging from 3.7% to 39.3% based on previous network studies (Peraza et al., 2015; van Wijk et al., 2010). Global and local efficiency were computed for each time window separately (Achard and Bullmore, 2007; Latora and Marchiori, 2001). Variability of efficiency was then assessed by integrating over all edge density thresholds and computing the standard deviation of the respective measure over time (Kim et al., 2017).

### 2.8. Statistical analysis

All statistical analyses were performed in R (https://www.R-project.org/). The variability of functional connectivity of each network and each connection was compared between the groups using a non-parametric multivariate ANOVA (MANOVA (Burchett et al., 2017)) with diagnosis as between-subject factor. The different k-means measures were also compared between the groups using non-parametric MANOVA followed by Kruskal-Wallis ANOVAs and post-hoc Dunn’s tests using false discovery rate (FDR) correction for multiple comparisons. Effect sizes for all group comparisons were calculated using r^2^ (see Supplementary Tables S3 and S5). Spearman’s rank correlations between dynamic functional connectivity measures that showed significant group differences and clinical variables in the DLB group were assessed for the CAF total score as a measure of cognitive fluctuation severity, the UPDRS motor subscale as a measure of the severity of Parkinsonism, the NPI hallucination subscale which was specifically scored for the severity and frequency of visual hallucinations, and the MMSE as a measure of global cognition. In the AD group, correlations with MMSE were calculated.

To assess the effect of dopaminergic medication on dynamic connectivity measures, DLB patients were divided into those patients who were taking dopaminergic medication and those who were not on these medications and all dynamic connectivity measures were compared between the two groups using Mann-Whitney U-tests (see Section 7 of the Supplementary Material).

Additionally, to investigate the possible influence of motion artefacts on the dynamic connectivity measures, we calculated correlations between mean framewise displacement and the dynamic connectivity measures across all participants (see Section 8 of the Supplementary Material).

## 3. Results

One AD patient had to be excluded due to coregistration errors. Additionally, two controls, six AD, and two DLB participants were excluded because of excessive motion. This resulted in 31 DLB, 29 AD patients, and 31 HCs for further analysis. The overall motion for all included participants did not differ between the groups (Kruskal-Wallis ANOVA; rotation, ¾=0.79, p=0.67; translation, ¾=0.67, p=0.71).

### 3.1 Demographics

Age and gender were not significantly different between the groups and the two dementia groups did not differ significantly in terms of overall cognition (MMSE and CAMCOG) and dementia duration (Table 1). As expected, the number of patients taking dopaminergic medication was significantly higher in the DLB group while the number of patients taking cholinesterase inhibitors was not different between the dementia groups. DLB patients had worse motor function, more visual hallucinations, and greater cognitive fluctuations than AD patients.

**Table 1:** Demographic and clinical variables, mean (standard deviation)

|  | HC (N=31) | AD (N=29) | DLB (N=31) | Between-group differences |
| --- | --- | --- | --- | --- |
| Male: female | 22:9 | 20:9 | 19:12 | $\chi^2=0.73$, $p=0.70^a$ |
| Study 1: study 2 | 15:16 | 13:16 | 12:19 | $\chi^2=0.60$, $p=0.74^a$ |
| Age | 76.4 (7.2) | 75.2 (8.6) | 78.1 (6.7) | $F_{2,88}=1.16$, $p=0.32^b$ |
| AChEI | - | 26 | 28 | $\chi^2=0.007$, $p=0.93^c$ |
| PD meds | - | 1 | 18 | $\chi^2=20.66$, $p<0.001^c$ |
| Duration | - | 3.7 (1.7)<sup>f</sup> | 3.4 (2.3) | $U=339$, $p=0.14^d$ |
| MMSE | 28.9 (1.1) | 21.8 (3.8) | 22.0 (4.3) | $t_{58}=0.20$, $p=0.85^e$ |
| CAMCOG | 96.7 (3.2) | 70.3 (13.5) | 73.3 (13.6) | $t_{58}=0.86$, $p=0.39^e$ |
| UPDRS III | 1.94 (2.8) | 3.5 (4.0) | 18.1 (10.0) | $t_{58}=7.32$, $p<0.001^e$ |
| CAF total | - | 1.00 (2.5)<sup>f</sup> | 4.8 (4.9)<sup>g</sup> | $t_{56}=3.66$, $p=0.001^e$ |
| NPI total | - | 5.9 (5.5)<sup>h</sup> | 14.6 (11.0)<sup>i</sup> | $t_{54}=3.68$, $p=0.001^e$ |
| NPI hall | - | 0<sup>j</sup> | 1.6 (1.8)<sup>i</sup> | $t_{53}=4.53$, $p<0.001^e$ |

AChEI, number of patients taking acetylcholinesterase inhibitors; AD, Alzheimer's disease; CAF total, Clinical Assessment of Fluctuations total score; CAMCOG, Cambridge Cognitive Examination; DLB, Dementia with Lewy bodies; Duration, duration of cognitive symptoms in years; HC, healthy controls; Mayo total, Mayo Fluctuations Scale; MMSE, Mini Mental State Examination; PD meds, number of patients taking dopaminergic medication for the management of Parkinson's disease symptoms; UPDRS III, Unified Parkinson's Disease Rating Scale III (motor subsection); NPI, Neuropsychiatric Inventory; NPI hall, NPI hallucination subscore <sup>a</sup> Chi-square test HC, AD, DLB; <sup>b</sup> One-way ANOVA HC, AD, DLB; <sup>c</sup> Chi-square test AD, DLB; <sup>d</sup> Mann Whitney U test AD, DLB; <sup>e</sup> Student's t-test AD, DLB.
<sup>f</sup> N=28; <sup>g</sup> N=30; <sup>h</sup> N=27; <sup>i</sup> N=29; <sup>j</sup> N=26.

### 3.2 Group differences in dynamic connectivity

The subject-specific values for the regularization parameter λ that resulted from the optimization procedure did not differ between the three groups (Kruksal-Wallis ANOVA, H_2_=0.06, p=0.97). Figure 3 shows matrices representing the mean SD of the strength of each RSN-to-RSN connection within each group. Overall mean variability of connectivity is shown in the bottom right panel of Figure 3. When considering average variability of each RSN, the overall MANOVA did not show a significant effect of diagnosis (F(10,442)=1.39, p=0.18). Similarly, when considering each individual RSN-to-RSN connection, the MANOVA did not reveal a significant group difference across all variables (F(96,4221)=1.02, p=0.43). Supplementary Table S3 and Supplementary Figure S2 show effect size estimates for all comparisons. Overall, effect sizes were largest for the comparison of HC and DLB participants while they were lowest for the comparison between both dementia groups.

**Figure 3:**
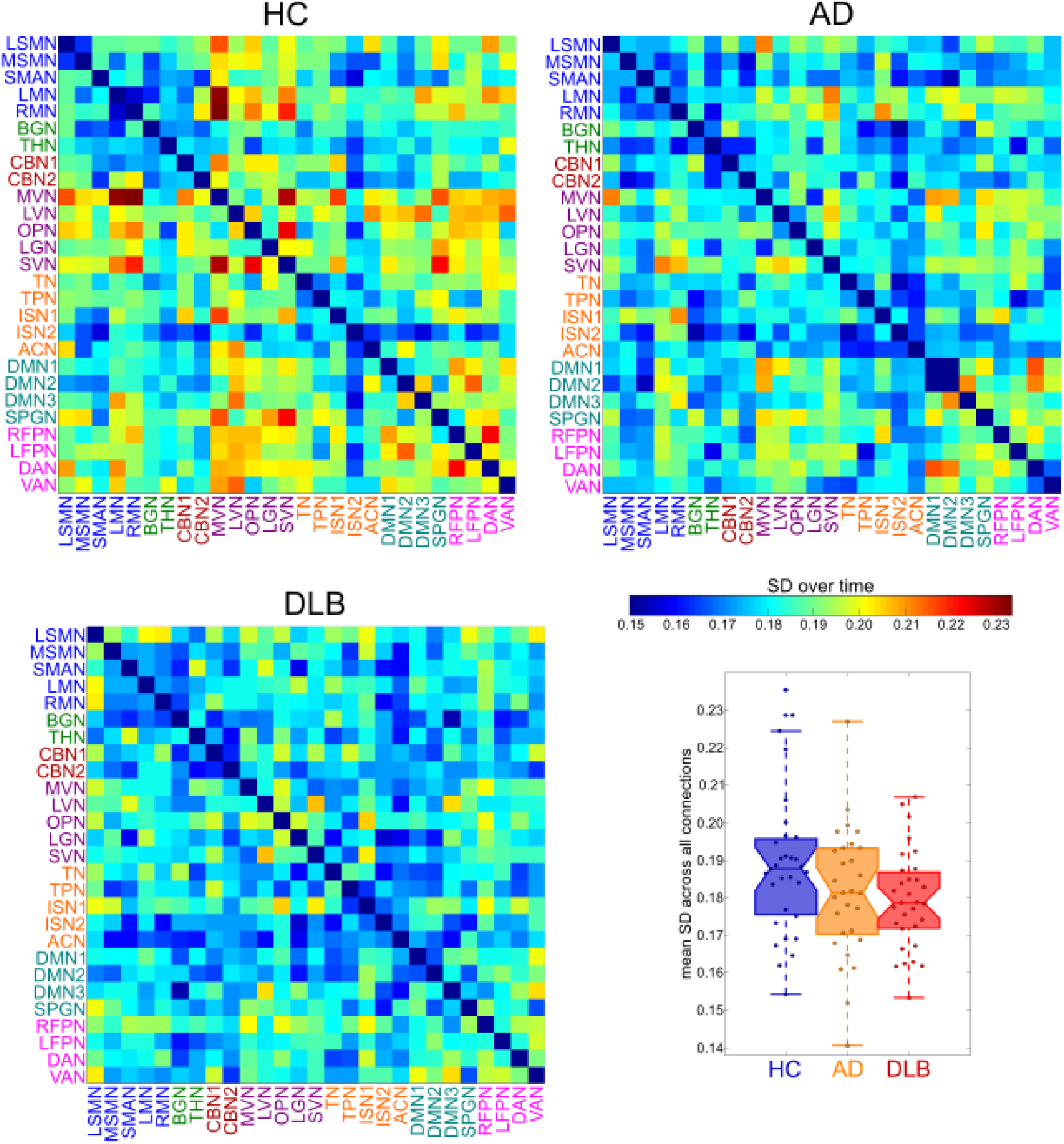
Results from dynamic functional connectivity analysis. Matrices representing mean standard deviation over time for all HC, AD, and DLB participants and boxplot showing a group comparison of mean standard deviation across all connections.

SD matrices were re-estimated using different window sizes from 18 to 28 TR showing that the overall appearance of the SD matrices was not dependent upon the specific choice of window size (Supplementary Figure S1).

### 3.3. K-means clustering

An optimal number of k=3 clusters was determined by the elbow criterion (Supplementary Figure S3). State 1 was characterized by relatively strong positive and negative between-network correlations (Figure 4). Especially strong positive correlations were present within the visual and the motor networks and between these two groups of networks (Figure 4E). Additionally, the motor and visual networks showed negative correlations with cognitive control, salience, and temporal networks and there was a strong connection between two components of the default mode network (DMN). In contrast, state 2 was characterized by much sparser connections, with weaker connectivity within visual and motor networks and a relative lack of connections between the two groups. There were a few positive connections between visual and default mode networks and additional positive connections between DMN and attention networks. State 2 was the most common state, being present in almost all participants and accounting for 50% of all time windows. Similar to state 2, state 3 was characterised by weaker connections and the relative absence of strong anti-correlations. In addition to some within-module connections in the visual, motor, and default mode networks, there were weak connections between visual and DMN and attention networks.

**Figure 4:**
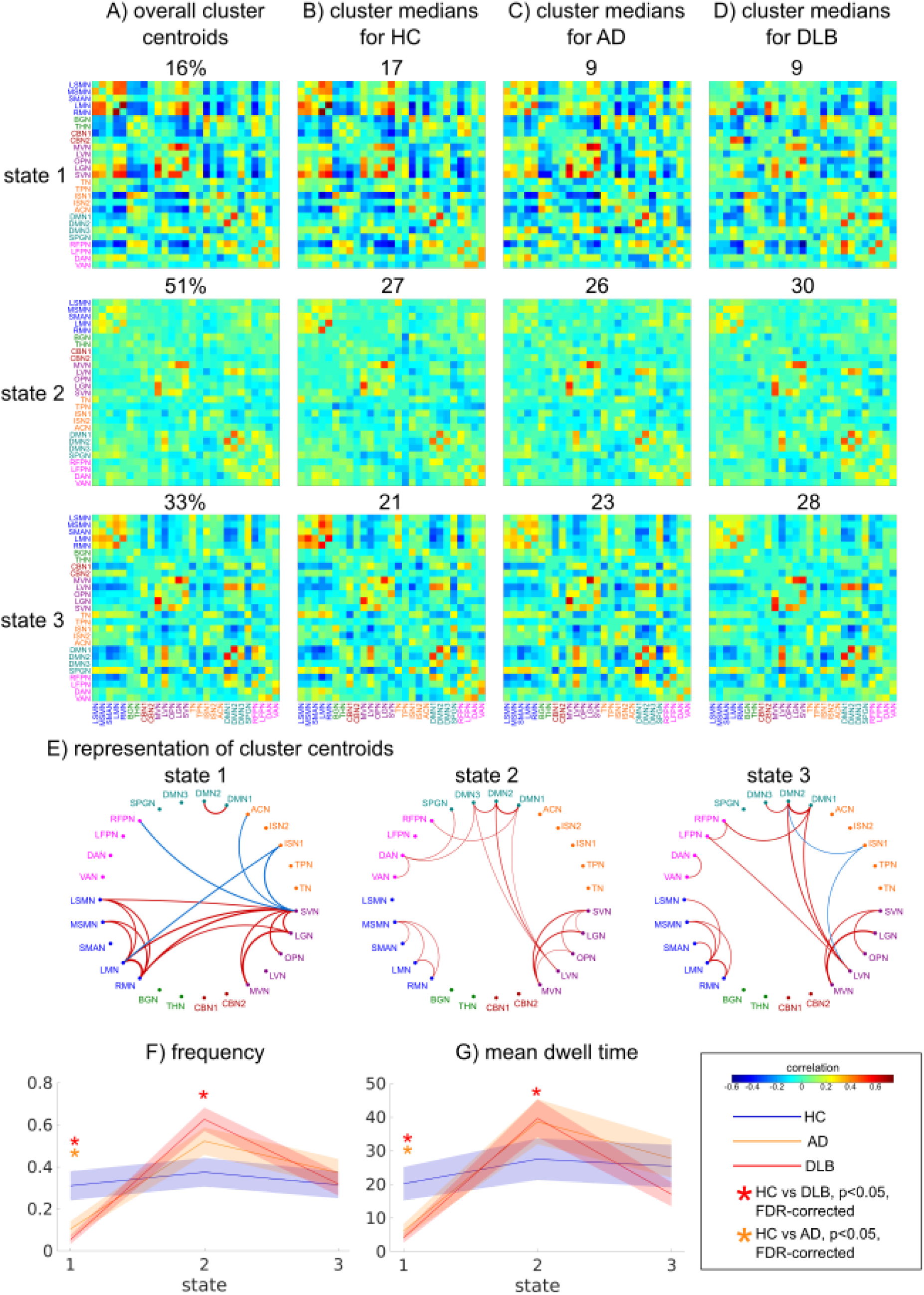
Results from k-means analysis. A) Centroids resulting from clustering on all windows with the overall percentage of windows assigned to the respective cluster. B) cluster medians in the healthy control (HC) group with the number of HC participants expressing this state. C) cluster medians in the Alzheimer’s disease (AD) group with the number of AD patients expressing this state. D) cluster medians in the DLB group with the number of DLB patients expressing this state. E) network representation of cluster centroids showing only the 5% strongest positive (red) and negative (blue) connections. F) comparison of frequency of occurrence between the three groups for each state, solid lines represent the means per group, shaded areas represent error bars of the standard error. G) comparison of mean dwell time in each state between the three groups. FDR-corrected p-values<0.05 are marked with an asterisk.

There were no significant differences between the groups in the number of state transitions or the intertransition interval (Supplementary Tables S4 and S5). The frequency of occurrence of the three states was not correlated with time, i.e. we did not observe an increase or decrease in the occurrence of any state over the duration of the scan (Supplementary Table S4).

Non-parametric MANOVAs revealed that there was a significant effect of diagnosis on frequency and mean dwell time across all three states (Supplementary Table S4 and S5). Follow-up univariate Kruskal-Wallis ANOVAs and pairwise post-hoc tests demonstrated that state 1 occurred less frequently in AD and DLB compared to controls with no difference between the dementia groups (Figure 4F and Supplementary Tables S4 and S5). In contrast, state 2 occurred more often in DLB compared to controls. However, there was no difference between HC and AD or between AD and DLB for state 2. The mean dwell time of state 1 and 2 followed the same pattern as the frequency, i.e. DLB patients spent shorter periods of time in state 1 and longer periods of time in state 2 than HC; AD patients spent shorter times in state 1 than HC with no difference for state 2, and there was no difference between the dementia groups (Figure 4G). There were no group differences in frequency or dwell time for state 3 (Supplementary Tables S4 and S5).

Several further analyses were performed to assess the robustness of the k-means analysis. Supplementary Figure S4 shows results for different numbers of clusters demonstrating that the main result of differences in frequency and dwell time of state 1 and 2 persisted when using a higher k. Additionally, repeating the k-means analysis with k=3 for different window sizes confirmed that the specific choice of window length did not influence the state identification (Supplementary Figure S5). We also performed split-half and bootstrap resampling which showed that states 1 and 2 were consistently identified in both split-half and all bootstrap resamples, while state 3 failed to be identified in some of the bootstrap resamples (Supplementary Figure S6).

### 3.4. Dynamic network measures

There was no difference between the groups in terms of variability of local efficiency (Figure 5B and Supplementary Table S4). In contrast, global efficiency differed significantly between the groups. Post-hoc tests revealed that it was less variable in DLB compared to controls with no significant difference between AD and HC as well as between the two dementia groups (Figure 5A and Supplementary Table S5). Figure 5C shows variability of global efficiency for different edge densities and indicates that the largest group difference occurred for edge densities of around 20%.

**Figure 5:**
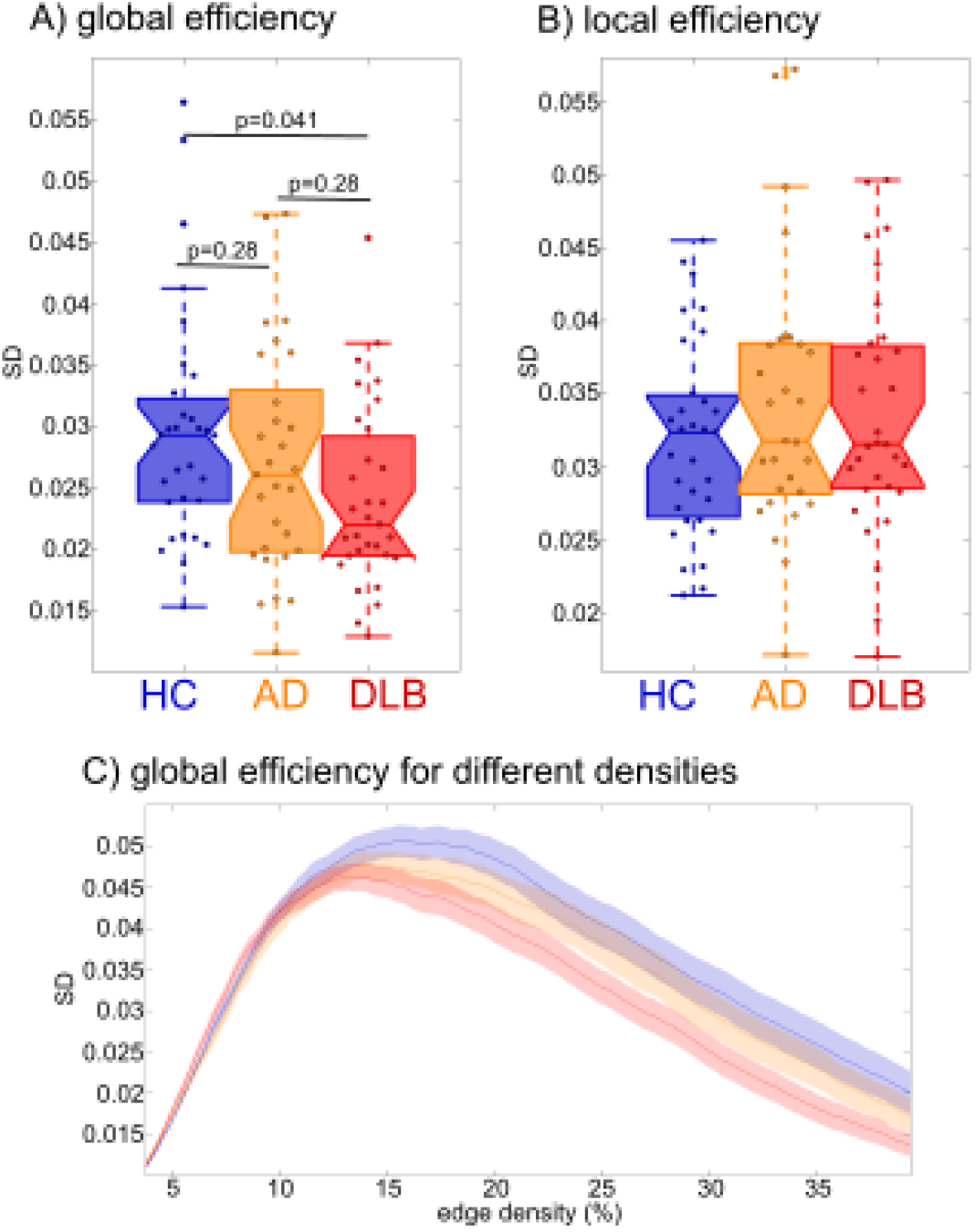
Results from dynamic network analysis. Comparison of the variability of A) global (p-values FDR-corrected) and B) local efficiency between groups. C) Variability of global efficiency for different edge densities.

### 3.5. Relation to clinical scores

Table S6 shows correlations between clinical variables and all dynamic connectivity measures that showed significant group differences in the DLB and AD groups separately. There were no significant correlations between cognitive fluctuation or visual hallucination severity and the dynamic connectivity measures. Frequency of state 2 was positively correlated with the UPDRS in DLB. However, this correlation did not survive correction for multiple comparisons. There were no significant clinical correlations in the AD group.

Comparing DLB patients who were on dopaminergic medication to those who were not, did not reveal any significant differences between the two groups (Supplementary Table S7).

There were no significant correlations between the dynamic connectivity measures that showed group differences and mean framewise displacement (Supplementary Table S8).

## 4. Discussion

In this study we investigated differences in functional connectivity dynamics and dynamic brain network topology between patients with DLB, patients with AD, and healthy controls. In terms of dynamic changes in overall network structure, we found reduced variability of global efficiency in the DLB group compared to controls which was not observed in the AD group. Using a state-based analysis it became evident that both dementia groups spent less time in a state of strong internetwork connectivity than controls. Additionally, DLB patients spent more time in a more sparsely connected state characterized by the relative loss of strong anti-correlations and an isolation of motor networks relative to other networks. While dynamic connectivity measures of the AD group were often between those of the control and DLB groups, we did not see significant differences in the direct comparison of both dementia groups.

### 4.1. State-based analysis

While the number of states visited and the number of state changes was not altered in the dementia groups, there was a significant difference in the type of state changes in dementia patients compared to controls. The frequency with which the control participants visited each of the three states was relatively balanced, i.e. they spent about a third of their time in each state. In contrast, the distribution of states in the AD and DLB groups was more out of balance with a clear decrease in frequency of state 1 in both dementia groups accompanied by an increased frequency of state 2 in DLB. In addition to visiting state 1 less often, the dementia patients also switched out of this state more rapidly and DLB patients stayed in state 2 for longer consecutive periods of time. In accordance with previous reports in healthy participants (Allen et al., 2014), development (Marusak et al., 2016), ageing (Viviano et al., 2017), and Parkinson’s disease (PD) (Kim et al., 2017), the most common state in the present study (state 2) was characterized by a sparse connectivity profile with relatively weak internetwork connections and the absence of strong anti-correlations. The frequency of this state has been linked to the amount of self-focussed thought (Marusak et al., 2016) and it has been suggested to represent a general connectivity pattern that participants spend most of their time in, with other states reflecting temporary deviations that might be due to cognitive, physiological, or motion-related processes (Viviano et al., 2017). State 1 deviated from this state by stronger positive and negative connections. It seems that AD as well as DLB patients remain in states of low inter-network connectivity and switch less often into more highly and specifically connected network configurations. This may relate to cognitive deficits associated with dementia in general even though we did not see any specific correlations between the time spent in different states and cognitive measures. A specific hallmark of state 1 is strong connectivity within visual and motor networks and between these two groups of networks that is not present in state 2. A reduced ability to switch into this state thus accords with Sourty *et al*. (2016) who found dynamic connectivity changes in DLB for networks related to visual processing using Hidden Markov Models (and thus is not directly comparable to our study). Another important characteristic of state 1 that differentiates it from state 2 is the existence of strong anti-correlations in the former. Anti-correlations between default mode and task-positive networks have been shown to be important for attentional function (Fox et al., 2005) and a loss of anti-correlations has been associated with aging, mild cognitive impairment, and cognitive impairment in PD (Baggio et al., 2015; Esposito et al., 2017). Our results further suggest that an absence of strong anti-correlations might also be a feature of more established neurodegenerative disease in the case of AD and DLB.

### 4.2 DLB-related changes in dynamic network topology

Regarding dynamic network topology, we found less variable global efficiency in DLB compared to controls. Global efficiency is a measure of communication efficiency across the whole brain network (Latora and Marchiori, 2001). In general, more pronounced variability of functional connectivity has been shown to be related to superior performance on a range of behavioral tests including attention and memory tasks (Jia et al., 2014) indicating that the dynamic and flexible engaging and disengaging of different brain regions seems to be crucial for efficient and adaptable communication within the brain (Zalesky et al., 2014). Reduced dynamics in turn can lead to less flexible, ineffective communication and a reduced ability of the network to respond to situational demands. The reduced variability of global efficiency in DLB might thus indicate a disease-related and abnormal rigidity of the brain network which might relate to the cognitive slowing (bradyphrenia) that is observed in DLB patients (Firbank et al., 2017). A previous static network analysis using the same study cohort found overall increased global efficiency in DLB (Peraza et al., 2015). However, our results suggest that this overall increase is in fact due to reduced variability indicating that DLB patients constantly stay in a network configuration of high global efficiency. In contrast, in healthy brains efficiency is temporally modulated which has been shown to represent more economical network dynamics allowing for a more specific response to situational demands (Zalesky et al., 2014). Similar to our results, Peraza *et al*. (2015) reported no difference between AD and controls with respect to global efficiency which indicates that static and dynamic changes in efficiency might be a specific feature of DLB that might not be associated with dementia *per se*. In contrast to our results, Kim *et al*. (2017) found increased variability of global efficiency in PD. However, this finding was not replicated in another study in PD patients with mild cognitive impairment (Díez-Cirarda et al., 2017) and thus further research will be needed to identify the specific changes related to these different Lewy body diseases.

### 4.3 Relation to clinical symptoms in DLB

Given the transient nature of clinical DLB symptoms such as visual hallucinations and cognitive fluctuations, we expected to find relations between symptom severity and dynamic connectivity measures. However, we did not observe any relation with respect to visual hallucinations and cognitive fluctuations, even before correcting for multiple comparisons. A possible reason for this might be the difference in time-scales: while our data only allowed the characterisation of dynamics during a 6-minute resting state scan, the time-scale of cognitive fluctuations and visual hallucinations can be on the order of minutes to hours and even days. Performing repeated scans with DLB patients at different times of the day or over several days might thus help to understand more about the relation between functional connectivity dynamics and clinical symptom severity. There was a trend for an association between frequency of state 2 and severity of Parkinsonism in DLB, i.e. an increased time spent in this sparsely connected state might relate to more severe Parkinsonism. Relative to state 1, this state was characterized by a disconnection of motor networks from other networks and the observed correlation might thus indicate that the isolation of motor networks might contribute to the severity of clinical motor symptoms. However, this is only an exploratory result that did not survive multiple comparison correction and further research will be needed to confirm this hypothesis.

### 4.4 Reliability of dynamic connectivity results

The interpretation, functional significance, and origin of dynamic functional connectivity have been the subject of an extensive debate (Hindriks et al., 2016; Laumann et al., 2017). However, recent studies using concurrent fMRI and EEG measurements point towards a neuronal origin of dynamic functional connectivity (Chang et al., 2013). Additionally, several studies have provided support for a cognitive role by showing that temporary changes in connectivity are related to changes in behavioural or vigilance states (Jia et al., 2014; Kucyi et al., 2017; Thompson et al., 2013) and cognitive performance in healthy older adults (Cabral et al., 2017). Finally, the study of dynamic functional connectivity in clinical populations has led to the identification of specific dynamic connectivity alterations associated with specific disorders which provides further evidence of the neurocognitive significance of time-varying functional connectivity (Damaraju et al., 2014; Jones et al., 2012; Sourty et al., 2016).

Although the sliding window approach has been widely applied to study dynamic functional connectivity (Allen et al., 2014; Damaraju et al., 2014; Hutchison and Morton, 2015; Jones et al., 2012; Marusak et al., 2016) its validity has been debated (Hindriks et al., 2016). Advantages are its interpretability and computational efficiency which make this kind of analysis especially suitable for the investigation of clinical questions. However, problematic aspects include the need for an *a priori*specification of parameters such as window length and the number of states for the k-means analysis and the possibility of spurious connectivity fluctuations which can arise due to noise sources such as head motion (Hutchison et al., 2013a). In the present study we applied several pre- and postprocessing steps to reduce the effect of these noise sources as far as possible (see Section 2.4). It was also ensured that the groups did not differ with respect to motion which makes it unlikely that the observed group differences were merely due to motion artefacts. Additionally, there was no significant relation between dynamic connectivity measures and mean framewise displacement indicating little influence of motion on the dynamic connectivity measures in our participants. Regarding the choice of window length, we showed that our results can be reproduced using windows of different lengths (see Supplementary Figures S1 and S5). While most previous studies examined a larger number of states (Allen et al., 2014; Damaraju et al., 2014; Hutchison and Morton, 2015; Marusak et al., 2016; Viviano et al., 2017), we focused on a smaller set of three states which was determined as the optimal number of states in our dataset and is comparable to a previous report in PD (Kim et al., 2017). The states tended to get more unstable as more states were added with states appearing that were specific to certain participants (Supplementary Figure S4). This might be due to the small number of participants and large heterogeneity in our sample. Nevertheless, we showed that the observed group differences in terms of frequency and dwell time remained largely unchanged for different values of k and states were reproducible on split-half and bootstrap resamples of the data which confirms the robustness of this approach. Notably, adding more states did not result in more significant group differences indicating that these three states represent the most important states in terms of dementia-related changes in connectivity dynamics.

### 4.5 Limitations

A limitation of this study is that over half of the DLB patients were on dopaminergic medication and scanned in the ON state which might have influenced their functional connectivity measures. However, dopaminergic medication has been shown to normalize connectivity towards more healthy levels (Tahmasian et al., 2015) suggesting that the observed group differences were not due to medication. Furthermore, we did not find differences in terms of dynamic connectivity measures between DLB patients who were taking dopaminergic medication compared to those not on these medications. All diagnoses were based on clinical assessment rather than pathological confirmation. However, the standardized diagnostic criteria that were used in this study have demonstrated high specificity when validated against autopsy findings (McKeith et al., 2000).

### 4.6 Conclusion

The loss of variability of global efficiency in DLB indicates an abnormally rigid brain network. This might be associated with less economical dynamics that can prevent specific and effective responses of the brain network to situational demands. This loss of dynamics was not observed in the AD patients and might therefore relate to clinical characteristics that are specific to DLB. However, the absence of correlations with visual hallucination and cognitive fluctuation severity indicates that contrary to our hypothesis and even though DLB is characterized by transient cognitive symptoms, their severity as measured by clinical scores might not directly relate to the dynamic changes in connectivity that are observable during a short resting state scan.

## 5. Acknowledgements

J.S. is supported by the Alzheimer’s Society Doctoral Training Centre at Newcastle University. M.K. is supported by the Engineering and Physical Sciences Research Council of the United Kingdom Grant EP/K026992/1. The research was supported by a Wellcome Trust Intermediate Clinical Fellowship (WT088441MA) to J.-P.T., Northumberland Tyne and Wear NHS Foundation Trust, by National Institute for Health Research (NIHR) Newcastle Biomedical Research Centre (BRC) based at Newcastle upon Tyne Hospitals NHS Foundation Trust and Newcastle University, and by Alzheimer’s Research UK.

